# Microgliosis driven by palmitate exposure alters energy metabolism and extracellular vesicles release that impact behavior and systemic metabolism

**DOI:** 10.1101/2024.04.04.587954

**Authors:** Gabriela C. De Paula, Blanca Aldana, Roberta Battistella, Rosalia Fernández Calle, Andreas Bjure, Iben Lundgaard, Tomas Deierborg, João M. N. Duarte

## Abstract

Dietary patterns that include an excess of foods rich in saturated fat are associated with brain dysfunction. Although microgliosis has been proposed to play a key role in the development of brain dysfunction in diet-induced obesity (DIO), neuroinflammation with cytokine over-expression is often not always observed. Thus, mechanisms by which microglia contribute to brain impairment in DIO are uncertain. Using the BV2 cell model, we investigated the gliosis profile of microglia exposed to palmitate (200 µmol/L), a saturated fatty acid abundant in high-fat diet and in the brain of obese individuals. We observed that microglia respond to a 24-hour palmitate exposure with increased proliferation, and with a metabolic network rearrangement that favors energy production from glycolysis rather than oxidative metabolism, despite stimulated mitochondria biogenesis. In addition, while palmitate did not induce increased cytokine expression, it modified the protein cargo of released extracellular vesicles (EVs). When administered intra-cerebroventricularly to mice, EVs from palmitate-exposed microglia *in vitro* led to memory impairment, depression-like behavior, and glucose intolerance, when compared to mice receiving EVs from vehicle-treated microglia. We conclude that microglia exposed to palmitate can mediate brain dysfunction through the cargo of shed EVs.

## Introduction

Excessive consumption of diets rich in saturated fat leads to obesity that, together with its metabolic complications and comorbidities, impacts the brain (Hartmann *et al*., 2020; Hoscheidt *et al*., 2022; Duarte, 2023). In animal models, metabolic syndrome upon diet-induced obesity (DIO) has been reported to trigger hippocampal metabolic alterations, synaptic dysfunction, and impairments in learning and memory processes (reviewed in García-Serrano & Duarte, 2020). Interestingly, just as in animal studies, a relatively short exposure to a high-fat and high-sugar diet (four days) is sufficient to deteriorate hippocampal-dependent learning and memory in healthy humans (Attuquayefio *et al*., 2017).

Obesity is associated with a state of low-grade inflammation and, in the brain, DIO triggers neuroinflammation and gliosis (Pistell *et al*., 2010; Thaler *et al*., 2012; Valdearcos *et al*., 2014, 2017; Cavaliere *et al*., 2019; de Paula *et al*., 2021). However, a controversy on neuroinflammatory processes induced by DIO can be found in the literature, such as a limited extension of gliosis and lack of cytokine overexpression in certain brain areas, such as hippocampus and cortex (Baufeld *et al*., 2016; Lizarbe, Cherix *et al*., 2019; Lizarbe, Soares *et al*., 2019; Mishra *et al*., 2019; Garcia-Serrano, Mohr *et al*., 2022; Skoug *et al*., 2024). Therefore, cellular communication mechanisms other than cytokine release must play a role in neuroinflammation in DIO, including the ability of microglia to shed eicosanoids or extracellular vesicles (EVs).

EVs are cell-derived membrane-surrounded vesicles (exosomes, ectosomes, microvesicles, apoptotic bodies, among others) that carry bioactive molecules (metabolites, nucleic acids and proteins) and, depending on their size, even cellular organelles. Thus, EVs participate in an orchestrated strategy of intercellular communication, including mediating inflammatory cues from microglia to other brain cells (reviewed in Caruso Bavisotto *et al*., 2019; Pascual *et al*., 2021). EVs are also transmitted across multiple bodily organs (Hu *et al*., 2023), although transfer across the blood-brain-barrier (BBB) might be restricted to particles of smaller size (Banks *et al*., 2020).

Palmitate, an abundant saturated fatty acid in diet, was found to increase in the cerebrospinal fluid of overweight and obese individuals relative to lean controls, and cerebrospinal palmitate concentration is correlated with body mass index and abdominal circumference, but not with obesity associated comorbidities, such as diabetes, dyslipidemia, or hypertension (Melo *et al*., 2020). Melo *et al*. further reported that intra-cerebroventricular (i.c.v.) injection of palmitate impairs synaptic plasticity and memory in mice, and increased astrocyte and microglia reactivity.

Using the BV2 microglia cell line, we investigated the gliosis profile and energy metabolism alterations induced by palmitate exposure, and tested the hypothesis that palmitate-exposed microglia utilize EVs as means of transmitting inflammatory messages to other cells.

## Methods

### Palmitate preparation

A stock solution of sodium palmitate was prepared by conjugation with bovine serum albumin (BSA). Briefly, 4.54 g of fatty acid-free BSA (#A7030, Sigma-Aldrich, St. Louis, MO-USA) was dissolved at 36°C in 16 mL of 150 mmol/L NaCl, and 61.2 mg of sodium palmitate (Sigma-Aldrich #P9767) was dissolved at 71°C in 4 mL of 150 mmol/L NaCl. The palmitate solution was then slowly added to the BSA solution while stirring, filtered with a sterile 22-µm polyvinylidene fluoride filter (Sigma-Aldrich #LGVV255F), and frozen at -20 °C in 11 mmol/L palmitate aliquots.

### BV2 cells

Murine BV2 microglial cells (#CRL-2469, ATCC, Manassas, VA-USA; RRID:CVCL_0182) were cultured in T75 polystyrene flasks (#83.3911, Sarstedt, Nümbrecht, Germany) at 37 °C with an atmosphere of 5% CO_2_, using Dulbecco’s modified Eagle’s medium (DMEM), containing 5 mmol/L glucose, 1 mmol/L pyruvic acid, and 4 mmol/L glutamine (#11885, Gibco, ThermoScientific, Göteborg, Sweden) supplemented with 10% fetal bovine serum (FBS, Gibco #10500064), 100 U/mL penicillin/streptomycin (Gibco #15140122). Cells were split every 2 days. For that, media was removed, cells were washed in Dulbecco’s phosphate-buffered saline without calcium and magnesium (DPBS, Gibco #14190250), and 1 mL of 0.05%(w/v) trypsin-EDTA with phenol red (Gibco #25300054) was added to detach the cells. After 5 minutes, 9 mL of fresh culture medium was added for re-seeding. For cell counting, suspended cells (10 μL) were mixed 1:1 with 0.4%(w/v) Trypan Blue (Sigma-Aldrich #T8154), the mixture was loaded in a hemocytometer, and live cells (unstained) were counted under a microscope.

For experiments, cells at passage 14-24 were seeded in polystyrene plates (unless otherwise stated) to a ∼40% confluence as described below for each method. After 6 hours, medium was replaced by DMEM without FBS for 12-16 hours before treatment with either with vehicle 0.25%(w/v) fatty acid-free BSA, or palmitate (PA, 200 µmol/L) in 0.25%(w/v) BSA for 24 hours. This palmitate concentration was selected from results of a dose-response pilot experiment, in which palmitate at 500 µmol/L during 24 hours induced a cell death rate above 50%. As a positive control of microglia reactivity, cells were incubated for 3 hours with 1 µg/mL lipopolysaccharides (LPS) from *Escherichia coli* O111:B4 (Sigma-Aldrich #L2630; Lot #000110081) in 0.25%(w/v) BSA.

### Cell proliferation, viability and apoptosis

Cellular proliferation rates were determined at given time points by counting cells as described above. The CyQuant MTT Cell Viability kit (#V13154, Invitrogen, ThermoScientific) and the Caspase-Glo 3/7 assay kit (#G8090, Promega, Nacka, Sweden) were used in 96-well plates (Sarstedt #82.1581.001) to determine cell viability and apoptosis, respectively, following the manufacturer’s instructions. Experiments were performed in quadruplicates with seeding at a density of 10^4^ cells/well.

### Oxygen consumption and proton efflux rates

Cellular oxygen consumption rate (OCR) and proton efflux measured as extracellular acidification rate (ECAR) were analyzed in the Seahorse XF96 (Agilent, Santa Clara, CA-USA) following the manufacturer’s instructions. Briefly, 10^4^ cells/well were seeded on the Seahorse cell culture plates using a total volume of 200 μL of culture medium. After 24 hours, cells were incubated with the treatment solutions (treatments are described in the figure legends).

*OCR assay*: Cells were washed and incubated for 1 hour with 180 μL assay medium [in mmol/L: 5 glucose (Agilent #103577-100), 1 pyruvate (Agilent #103578-100), 2 glutamine (Agilent #103579-100) in XF DMEM medium (Agilent #103575-100)] in atmospheric air at 37°C. After equipment calibration, baseline respiration measurements were followed by 1.5 μmol/L oligomycin addition to determine ATP-linked and proton leak-driven respiration. The mitochondrial uncoupler FCCP (carbonyl cyanide-p-trifluoromethoxy-phenylhydrazone, 0.5 μmol/L) was added to induce maximal respiratory capacity. Non-mitochondrial respiration was determined after the addition of 0.5 μmol/L rotenone plus 0.5 μmol/L antimycin A (inhibitors of complex I and complex III, respectively).

*ECAR assay*: Cells were incubated in assay medium without glucose or pyruvate (sodium bicarbonate and FBS were absent) in atmospheric air at 37 °C. Baseline ECAR was measured after addition of 10 mmol/L glucose. The conversion of glucose to pyruvate, and production of lactate that is released with protons leads to medium acidification that can be used as surrogate of glycolysis. Oligomycin (1 μmol/L) was added to inhibit mitochondrial ATP production, revealing the cellular maximum glycolytic capacity. Non-glycolytic ECAR was measured after addition of the glycolysis inhibitor 2-deoxy-D-glucose (2-DG, 50 mmol/L).

OCR and ECAR values were normalized to total protein content, as determined with the bicinchoninic acid assay (kit from Pierce, #23227, ThermoScientific).

### Quantitative real-time polymerase chain reaction (qPCR)

Cells were seeded onto 24-well plates (Sarstedt #83.3922.005) at a density of 5x10^4^ cells/well. After treatments, cells were washed with ice-cold phosphate-buffered saline (PBS, Gibco #18912014), treated with 100 μL trypsin as above, and re-suspended in 400 μL of culture medium. Total RNA was extracted with TRIzol (Invitrogen #15596026) following the manufacturer’s instructions, and RNA samples (500 ng) were reverse transcribed using the qScript cDNA Synthesis Kit (#95047, QuantaBio, England). The resulting cDNA was used for qPCR with the reaction cocktail PerfeCTa SYBR Green FastMix (QuantaBio #95072), and the primer pairs in table 1. Reactions were run in duplicates, and relative gene expression levels were calculated with the ΔΔCT method using L14 as internal control gene.

**Table 1.**
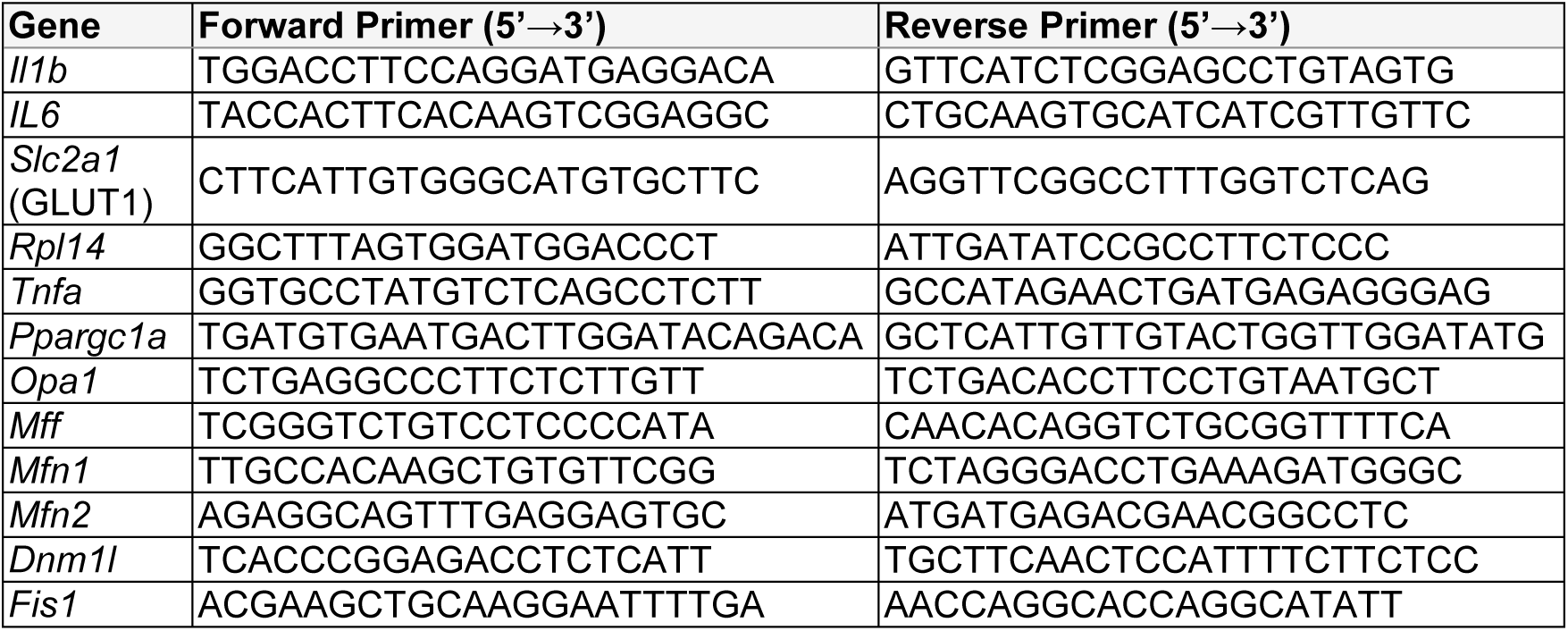
Nucleotide sequence of primers used for real time-PCR.

### TNF-α ELISA

Media were centrifuged at 2,000 x g and 4°C during 10 minutes to remove debris, and the supernatant was saved at -20°C until use. Concentration of TNF-α was determined with a Mouse TNF-α ELISA kit (Abcam, #ab208348).

### Immunoblotting

Cells were seeded onto 6-well plates (Sarstedt #83.3920.005) at a density of 3x10^5^ cells/well, and treated as described above. Cells were washed with PBS, and proteins were extracted with a lysis buffer [in mmol/L: 150 NaCl, 1 ethylenediaminetetraacetic acid (EDTA), 50 tris(hydroxymethyl)aminomethane (Tris)-HCl, 1% (v/v) Triton X-100, 0.5% (w/v) sodium deoxycholate, 0.5% (w/v) sodium dodecylsulfate (SDS), pH 8.0] containing cOmplete protease inhibitors (Roche, Switzerland). After protein determination, we separated 30 µg of protein by SDS-PAGE, and transferred onto nitrocellulose membranes as described elsewhere (Skoug *et al*., 2022). Membranes were blocked for 1 hour at room temperature with 5%(w/v) BSA in Tris-buffered saline (in mmol/l: 20 Tris, 150 NaCl, and pH 7.6) containing 1% Tween-20 (TBS-T), incubated overnight at 4 °C with the OxPhos Complexes antibody cocktail (Invitrogen #45-8099), diluted 1:1250 in blocking solution. Membranes were then washed three times in TBS-T for 15 minutes, and incubated for 2 hours at room temperature with horseradish peroxidase-conjugated anti-mouse IgG secondary antibody (dilution 1:5000, Abcam #ab6789). After washing again, immunoblots were developed on the ChemiDoc (Bio-Rad, Sundbyberg, Sweden) using the SuperSignal West Pico PLUS Chemiluminescent substrate (ThermoScientific #34580). For each independent experiment, the 3 experimental groups were run in parallel in the same gel, and signal from each band was normalized to the average of that in the 3 groups.

### Mitotracker staining and immunofluorescence microscopy

Cells (5x10^4^) were seeded onto a 35-mm glass-bottom dish coated with poly-D-lysine (#P35GC-1.5-14-C, MatTek, Ashland, MA-USA), and treated as described above. Then, cells were incubated for 30 minutes at 37°C with 100 nmol/L of MitoTracker (Invitrogen #M7512; 1 mmol/L stock solution in DMSO). Stained cells were rinsed in PBS, and fixed with buffered 4% formaldehyde (Histolab, Askim, Sweden) for 15 minutes at room temperature. Fixed cells were rinsed 3 times, permeabilized during 10 minutes with 0.2%(v/v) Triton X-100 in PBS, blocked for 30 minutes in 5%(v/v) goat serum and 0.3%(v/v) Triton X-100 in PBS, and incubated overnight at 4 °C with blocking solution containing the primary antibodies rabbit anti-allograft inflammatory factor 1 (Iba1, dilution 1:200; #019-19741, Fujifilm Wako, Japan) and mouse anti-β-actin (dilution 1:400, Sigma-Aldrich #A3854). Cells were then incubated for 1 hour with the secondary antibodies AF488-conjugated anti-rabbit IgG (dilution 1:500, #AB150077, Abcam, Cambridge, UK) and AF647-conjugated anti-mouse IgG (dilution 1:500, Invitrogen #A21235), washed in PBS, mounted with 4,6-diamidino-2-phenylindole (DAPI)-containing Fluoroshield (Sigma-Aldrich #F6057), and imaged in a Nikon A1RHD confocal microscope with a CFI Apochromat TIRF 100 × Oil, NA 1.49 (Nikon Instruments, Tokyo, Japan). Images were acquired with NIS-elements (Laboratory Imaging, Nikon), and analyzed in ImageJ (NIH, Bethesda, MD-USA).

### 13C tracing experiments

Cells (2.2 x 10^6^) were seeded onto 10-cm dishes (#89089-612, ThermoScientific) and treated as above, using glucose-free DMEM (#11966025, ThermoScientific) supplemented with 5 mmol/L [1-^13^C]glucose (99% C13 atom, Cortecnet, Voisins le Bretonmeux, France). [1-^13^C]glucose was present during the 24 hours of treatment with palmitate or vehicle, or for 21 hours ahead and during the 3 hours of LPS treatments. Cells were washed with ice-cold DPBS and frozen in N_2_ (l), and metabolites were extracted with 1 mL 80%(v/v) methanol. Then, samples were sonicated for 30 minutes at 4 °C, and centrifuged at 13,000 x *g*, 4℃, for 30 minutes. Supernatants were dried using a Savant SpeedVac (ThermoScientific) operating at room temperature.

Dried samples were re-suspended in 100 mmol/L sodium phosphate buffer pH 7.4 prepared in ^2^H_2_O (>99.9%, Sigma-Aldrich), containing 0.01% NaN_3_. Sodium fumarate (0.3 µmol) was added as internal standard, and samples were transferred into 5 mm Wilmad NMR tubes (Sigma-Aldrich).

NMR spectra were acquired on a Avance III HD 600 MHz spectrometer with a standard TCI cryoprobe (Bruker Nordic, Solna, Sweden). Solvent-suppressed ^1^H-NMR spectra were acquired with the ZGPR pre-saturation pulse sequence with spectral width of 9 kHz, 3 s acquisition time, a relaxation delay of 22 s, and 24 scans per cell extract. ^1^H-decoupled ^13^C-NMR spectra were acquired using the ZGPG30 sequence with 30 kHz spectral width, 2 s acquisition time, and a relaxation delay of 2 s. To achieve adequate signal-to-noise ratio, ^13^C spectra were recorded with at least 30,000 scans. Spectra were analyzed by line fitting using NUTS (Acorn NMR, Fremont, USA), as in previous studies (*e.g.* Duarte *et al*., 2007). Multiplet fractions from aliphatic carbons of glutamate were used in tcaCALC running on MATLAB 2019a (MathWorks, Natick, MA-USA) to determine rates (relative to citrate synthase) of pyruvate dehydrogenase (PDH), anaplerosis through pyruvate carboxylase (Y_PC_), anaplerosis through other pathways such as propionate or glutamine (Y_S_), and pyruvate kinase (PK) (Alger *et al*., 2021).

### EVs isolation

Cell media were collected and centrifuged at 400 x *g* for 5 minutes at room temperature to remove any cells. The obtained supernatant was centrifuged at 2,000 x *g* at 4 °C for 10 minutes to remove cell debris, and the then at 30,000 x g at 4 °C for 30 minutes to remove large particles such as apoptotic cell bodies. The supernatant was again ultracentrifuged at 100,000 x *g* at 4 °C for 70 minutes to pellet the EVs. The resulting pellets were gently re-suspended in 20 μL of DPBS, and either kept at 4 °C overnight (for NTA and injection into the mouse brain) or stored at -20 °C (for proteomics).

### Nanoparticle tracking analysis (NTA)

EVs were analyzed using the NanoSight LM10 (Malvern Panalytical, Malvern, UK). Samples were diluted in PBS to 10^6^-10^9^ particles/mL, and injected at 50 μL/min and at room temperature (21-22 °C). Particles were tracked 5 times during 60 s, and analyzed with NanoSight NTA 3.4 (Malvern Panalytical) to determine particle diameter.

### Mass spectrometry (MS) for proteomics

EVs (20 µL) were mixed with 30 µL of RIPA buffer (Sigma-Aldrich #R0278), and sonicated with 30 cycles of 15 s ON-OFF using a BioRuptor (Diagenode, Denville, NJ-USA). The EV lysate was reduced with 10 mmol/L dithiothreitol at 56 °C for 30 minutes, followed by alkylation with 20 mmol/L iodoacetic acid for 30 minutes in the dark. Samples were precipitated with ice-cold ethanol 90%(v/v) overnight at -20 °C. Samples were centrifuged at 14,000 x g for 10 minutes. The pellets were air-dried, re-dissolved in 50 µL 100 mmol/L ammonium bicarbonate, sonicated, and centrifuged again. Supernatants were collected, and protein concentration was determined using a DeNovix nanospectrophotometer (AH diagnostics, Solna, Sweden). Protein samples (15 µg) were digested overnight at 37 °C with trypsin (Promega, Madison, WI-USA) in a protein:trypsin ratio of 50:1 (w/w). The digestion was stopped by 5 µL 10%(v/v) trifluoroacetic acid. Samples were dried using a SpeedVac, and re-dissolved in a mixture of 2%(v/v) acetonitrile and 0.1%(v/v) trifluoroacetic acid.

Samples were analyzed in an Orbitrap Eclipse Tribrid mass spectrometer coupled with an Ultimate 3000 RSLCnano system (ThermoFischer). The HPLC used a two-column setup: peptides were loaded into an Acclaim PepMap 100 C18 pre-column (75 μm x 2 cm; ThermoFischer) and then separated with the flow rate 300 nL/min in an EASYspray column (75 μm x 25 cm, nanoViper, C18, 2 μm, 100 Å; ThermoFischer). The column temperature was set 45 °C. Peptides were eluted with a nonlinear gradient using 0.1%(v/v) formic acid in water as solvent A, and 0.1%(v/v) formic acid in 80%(v/v) acetonitrile as solvent B. Solvent B was maintained at 2% during 4 minutes, increased to 25% during 100 minutes, to 40% during 20 minutes, to 95% during 1 minute, and finally kept at 95% for 5 min to wash the column.

Samples were analyzed with the positive data-dependent acquisition (DDA) mode. The full MS resolution was set to 120,000 at normal mass range, and the automatic gain control target (AGC) was set to standard with the maximum injection time to auto. The full mass range was set 350-1400 m/z. Precursors were isolated with the isolation window of 1.6 m/z and fragmented by HCD with the normalized collision energy of 30. MS^2^ was detected in the Orbitrap with the resolution of 15,000, and AGC and maximum injection time were set to standard and 50 ms, respectively.

The raw DDA data were analyzed with Proteome Discoverer 2.5 Software (ThermoScientific), and the peptides were identified using SEQUEST HT against the UniProtKB Mouse database (UP000000589) with the following parameters applied: cysteine carbamidomethylation as static modification, and *N*-terminal acetylation and methionine oxidation as dynamic modification. Precursor tolerance was set to 10 ppm, and fragment tolerance was set to 0.05 ppm. Up to 2 missed cleavages were allowed. Percolator false discovery rate (FDR) was used for peptide validation at a q-value below 0.01. The extracted chromatographic intensities were used to compare peptide abundance across samples.

### Proteome analysis

We adopted an all *versus* all contrast approach for the analysis of the protein signal intensities. Only proteins that were present in at least 2/3 independent experiments of each group were used for further analysis. A quantile-based regression was used to replace missing values with random draws. R version 3.6.2 (RRID:SCR_001905) was used for principal component analysis (PCA), and for differential enrichment testing using the package DEP: Differential Enrichment analysis of Proteomics data (https://rdrr.io/bioc/DEP). Significance thresholds were set to adjusted α=0.05 and log_2_ of the fold change=1. Significant findings were used for gene ontology analysis in using ShinyGO 0.77 (http://bioinformatics.sdstate.edu/go) and the Reactome pathway database.

### Animals

Experiments on mice adhered to the EU Directive 2010/63/EU, were approved by the Malmö/Lund Committee for Animal Experiment Ethics (permit #5123/2021), and are reported following the ARRIVE guidelines (Animal Research: Reporting In Vivo Experiments, NC3Rs initiative, UK). Sixteen 8-week-old male Swiss mice were obtained from Janvier Labs (Le Genest-Saint-Isle, France) and housed in groups of 4 under controlled conditions of humidity (55-60%) and temperature (21-23 ^◦^C) with a 12-hour light:dark cycle (lights on at 7:00). Chow and water were provided *ab libitum*. Mice were randomly selected (by coin tossing) to receive EVs either palmitate- or vehicle-treated BV2 cells, so that each cage housed 2 mice from either experimental group (total n=8/group). Body weight was evaluated at the start and end of the study.

### Injection of EVs

Mice were anesthetized with isoflurane (induction with 5%; maintenance with 2-3%), and their heads were fixed at a 45° angle in a stereotactic frame (Kopf Instruments, Tujunga, CA-USA) on a heated pad. After craniotomy under magnification, a glass micropipette (diameter 20-40 μm) was used to deliver 500 ng protein from fresh BV2-derived EVs to the lateral ventricle (anterior/posterior, 0.34 mm from bregma; medial/lateral, 1.0; dorsal/ventral, -2.2 mm from the skull; injection site confirmed in pilot experiments injecting trypan blue). EVs were delivered in multiple microinjections over 3 minutes using air pressure from a PLI-100A Pico-Injector (Harvard Apparatus, Cambridge, UK), and were allowed to diffuse during 4 minutes before the needle was withdrawn. After, animals received subcutaneous saline for hydration (1 mL), and 5 mg/kg Bupivacaine (Marcain, Aspen Nordic, Ballerup, Danmark) for pain relief upon recovery.

### Glucose tolerance test (GTT)

Food was removed for 5 hours and, thereafter, blood glucose was measured from tail tip blood with the Accu-Chek Aviva glucometer (Roche, Manheim, Germany). Then, mice were given 2 g/kg glucose i.p. from a 30% (w/v) solution in saline, and blood glucose was measured from the tail tip at 15, 30, 60, 90 and 120 minutes after injection.

### Behavior

Mice were tested between 8:00 and 17:00, in a cubic arena with side length of 50 cm, with room light adjusted to an illuminance of 15 lux. All experiments were recorded by an infrared camera into AnyMaze 6.0.1 (Stoelting, Dublin, Ireland). Mice were habituated to the experimental setup by exploring the empty arena during 5 minutes in 3 consecutive days. In the last habituation session, arena exploration was analyzed for total number of crossings between quadrants, time spent in center or perimeter (delineated at 8 cm from the walls), number of rearing events, and the clockwise and anti-clockwise rotations.

In the following day, memory performance was assessed with novel object recognition (NOR) and novel location recognition (NLR) tasks, as detailed previously (Garcia-Serrano, Mohr *et al*., 2022). Briefly, mice were allowed to explore the arena for 5 minutes with two identical objects (familiarization phase). The mice returned to their home cage for 1 hour (retention phase), and after were reintroduced for 5 minutes with one of the objects either replaced by a novel object or relocated in space (retention phase). The time spent exploring each object was analyzed in both familiarization and recognition phases.

The sucrose-plash test was used to evaluate depression-related behavior. Mice were sprayed with ∼1 mL of a 10% sucrose solution on their dorsal coat, and analyzed for latency between spraying and first grooming event, and total duration of grooming during 5 minutes. (de Paula *et al*., 2021)

### Immunofluorescence microscopy in brain slices

Animals under isoflurane anesthesia were transcardially perfused with ice-cold PBS, followed by 4% formaldehyde. Fixed brains were removed, embedded in 4% formaldehyde for 24 h, and cryoprotected in a 30% sucrose solution in PBS at 4 °C. Free-floating coronal sections (30 µm) were blocked for 2 hours at room temperature with 5%(v/v) goat serum solution in PBS containing 0.3%(v/v) Triton X-100, and then incubated overnight at 4 °C with primary antibodies anti-Iba1 (1:200), and with anti-glial fibrillary acidic protein (GFAP) pre-tagged with AF488 (1:500; ThermoScientific #53-9892-82). After washing with PBS, sections were incubated with the AF568-conjugated goat anti-rabbit IgG antibody (1:500, ThermoScientific #A-21069) for 2 hours at room temperature, washed again, and incubated with 4,6-diamidino-2-phenylindole (DAPI; 1 µg/mL in PBS; ThermoScientific #62247) for 10 minutes. Slices were then mounted onto slides with ProLong Glass antifade medium (Invitrogen #P36980), and imaged in a A1RHD confocal microscope interfaced with NIS-elements (Nikon). Z-stack-projected images were processed and analyzed in ImageJ for area of Iba1 and GFAP immunoreactivity as previously described (Skoug *et al*., 2024). Morphology of Iba1^+^ cells (3 cells/region/mouse) was analyzed with the AnalyzeSkeleton plugin (Arganda-Carreras *et al*., 2010) to extract the number of cell processes and their length, and the number of branching points.

### Statistical analysis

All data were analyzed using Prism 10.2.0 (GraphPad, San Diego, CA-US). Unless otherwise stated, data are presented as mean±SD of n independent experiments. Normality was assessed with the Kolmogorov-Smirnov test, or the Shapiro-Wilk test for small sample sizes. Data not showing a Gaussian distribution were either analyzed with non-parametric tests, or log-transformed before ANOVA. Behaviour results deviating from a normal distribution are represented in boxplots extending from the 25^th^ to 75^th^ percentiles, line at median, and whiskers to the minimum and maximum values. Two-group comparisons were made with 2-tailed Student’s t-tests or Mann Whitney tests. Multiple groups were analyzed with either a Kruskal-Wallis test followed by Dunn’s multiple comparisons or with ANOVA followed by *post hoc* comparisons using the Holm-Šídák method. Significance was accepted for P<0.05.

## Results

### Palmitate induces gliosis without cytokine overexpression

BV2 cells responded to 200 µmol/L palmitate exposure during 24 hours with increased proliferation, but at a slower rate than with LPS treatment (Figure 1A), which was confirmed by an increased cell viability in the MTT assay (+66% for palmitate; +60% for LPS; Figure 1B). There was also increased caspase activity in palmitate- and LPS-treated cells relative to vehicle (+19% for palmitate; +26% for LPS; Figure 1C).

**Figure 1.**
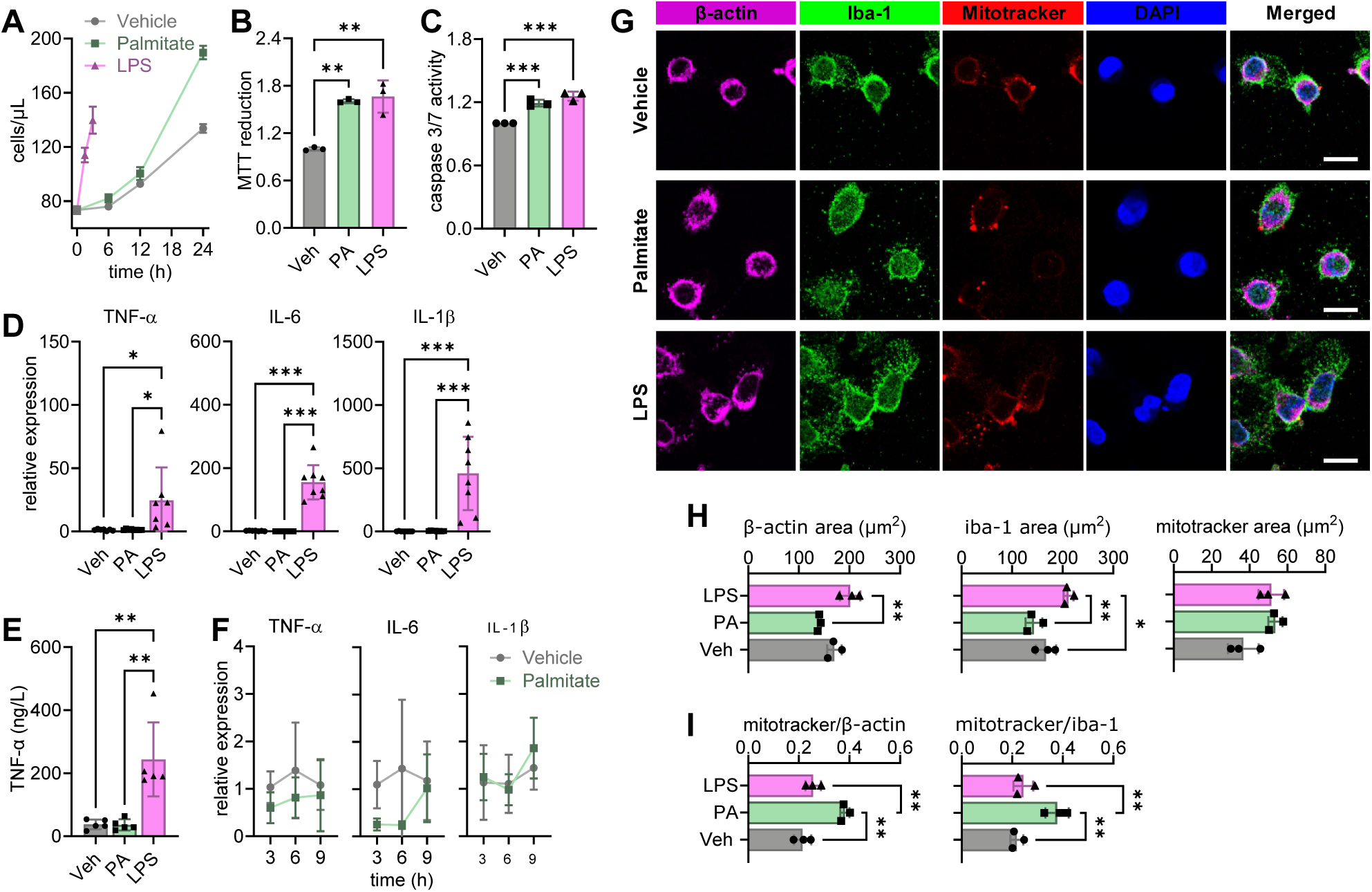
Palmitate exposure induces gliosis without exacerbated cytokine production, and increases mitochondria content. BV2 cells were incubated with vehicle (Veh), 200 µmol/L palmitate (PA) for 24 hours or 1 µg/mL LPS for 3 hours. (A) Cell counts before, and after 6, 12 and 24 hours of treatment, or after 1.5 and 3 hours for LPS. (B) Relative cell viability measured by MTT reduction. (C) Relative activity of caspase 3/7. (D) Relative expression of TNF-α, IL-6 and IL-1β, and (E) concentration of TNF-α in the medium after treatment. (F) Relative expression of cytokines during the initial 9 hours of palmitate exposure. (G) Representative immunofluorescence micrographs for mitotracker, β-actin and Iba1 (scale bar is 10 µm). (H) Mean area occupied by mitotracker, β-actin and Iba1 signal per cell, and (I) area ratios of mitotracker to β-actin and to Iba1. Cells were analyzed within 3-4 fields of view from 3 independent experiments. Data is shown as mean±SD of 3-8 independent experiments, represented by the individual symbols. *P<0.05, **P<0.01, ***P<0.001 depict differences in comparisons following significant effects in ANOVA.

Expression of the cytokines TNF-α, IL-1β and IL-6, and levels of TNF-α released into the medium were unaffected by palmitate treatment, while significantly increasing upon LPS exposure, relative to vehicle-treated cells (Figure 1D-E). We tested whether expression of cytokines was transiently modified at shorter palmitate exposure periods. Palmitate failed to increase cytokine expression at any of the time points assessed up to 9 hours of exposure (Figure 1F). Then, we set to determine whether palmitate expanded the area of the cells and their mitochondria, since palmitate can activate mitochondria in other cell models (Egnatchik *et al*., 2014). When compared to vehicle, palmitate-treated cells were of similar size, while LPS-treated cells increased the cell area as assessed by the immunoreactivity to either β-actin or the microglia marker Iba1 (Figure 1G-H). Interestingly, the area of mitotracker staining tended to be increased by both palmitate (P=0.054) and LPS (P=0.061) relative to vehicle (Figure 1G-H), and palmitate but not LPS exposure resulted in increased fraction of mitotracker fluorescence area relative to total cell area depicted by either β-acting or Iba1 (respectively, +78% and +73%; Figure 1I).

Altogether, these findings suggest that BV2 microglia exposed to palmitate respond with increased proliferation and expanded mitochondrial density, but not with a typical neuroinflammatory response that involves cytokine overexpression.

### Distinct metabolism alterations after palmitate and LPS exposure

Quiescent microglia mainly rely on oxidative phosphorylation for ATP production (Bernhart *et al*., 2010; Won *et al*., 2012). Upon activation, microglia stimulate glucose uptake and glycolysis to meet enhanced energy demands (Ghosh *et al*., 2018; Gimeno-Bayón *et al*., 2014). Thus, we set to measure palmitate-induced alterations of mitochondrial respiration. When compared to vehicle, cells exposed to either palmitate or LPS showed significantly lower baseline respiration, ATP-associated respiration, maximal respiration, and spare capacity (Figure 2A-C). Proton leakage and non-mitochondrial oxygen consumption were similar across all groups (Figure 2C). Since palmitate lowered mitochondrial respiration capacity, despite increased mitochondrial density, we then measured the density of mitochondrial respiration complexes and ATP synthase. Relative to vehicle-treated cells, palmitate but not LPS significantly increased the density of complexes I, II and IV by 57%, 63% and 73%, respectively (Figure 2D-E). We further determined the expression of genes involved in mitochondrial dynamics, and observed that both palmitate and LPS induced a significant increase in the expression of PGC-1α (Figure 1F), which is a key regulator of mitochondria biogenesis (Halling & Pilegaard, 2020). In turn, the expression of genes involved mitochondrial quality control by fusion (Opa1, Mfn1/2) and fission (Mff, Drp1, Fis1) was similar between the three groups (Figure 1F).

**Figure 2.**
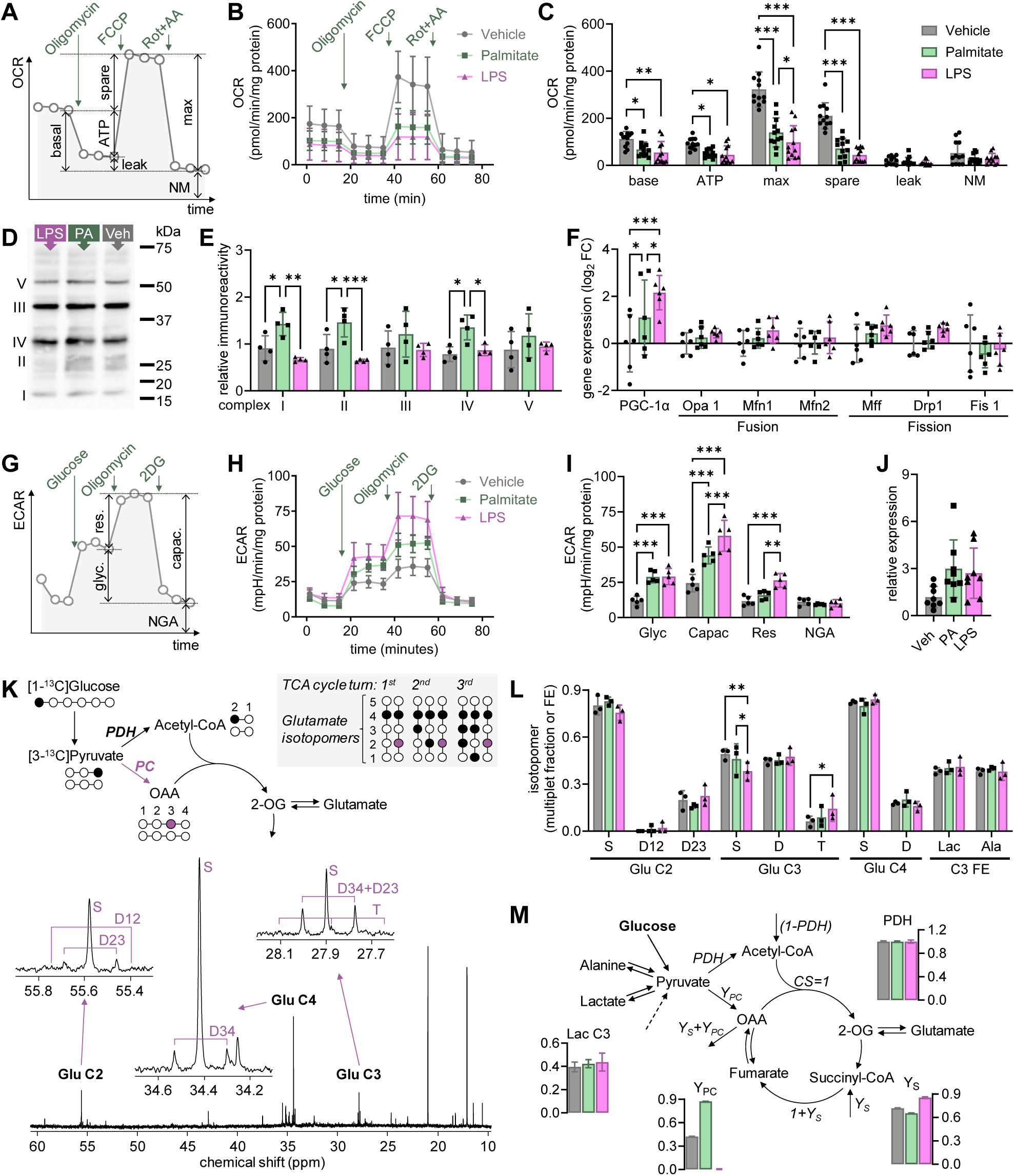
Palmitate exposure modulates energy metabolism in BV2 cells. BV2 cells were incubated with vehicle (Veh), 200 µmol/L palmitate (PA) for 24 hours or 1 µg/mL LPS for 3 hours. (A) Schematic representation of experiments for OCR measurements depicting the calculated parameters upon addition of oligomycin (1.5 mmol/L), FCCP (0.5 mmol/L), and antimycin A (0.5 mmol/L) plus rotenone (0.5 mmol/L): basal respiration (basal), proton leak-driven respiration (leak), ATP synthesis-linked respiration (ATP), maximal respiration capacity (max), spare respiration capacity (spare), and non-mitochondrial oxygen consumption (NM). (B-C) oxygen consumption rate (OCR) measured for 3 cycles within each respiration state (B), and calculated respiration parameters (C). (D) Representative immunoblotting experiment against the four complexes of the electron transport chain and ATP synthase (complex V), after separation of 30 µg of protein by SDS-PAGE. (E) Relative immunoreactivity signal from the 5 complexes in 4 independent experiments. For a given protein, signal within each band was normalized to the average of that in the 3 experimental groups. (F) Expression of genes involved in mitochondria biogenesis, fusion and fission. (G) Schematic representation of experiments for ECAR measurements depicting the calculated parameters upon addition of oligomycin (1 mmol/L) and 2-deoxy-D-glucose (2DG, 50 mmol/L): basal glycolysis (glyc), glycolytic reserve (res), glycolytic capacity (capac), and non-glycolytic medium acidicitation (NGA). (H-I) extracellular medium acidification rate (ECAR) measured for 3 cycles within each respiration state, and calculated glycolytic parameters. (J) Relative expression of *Slc2a1* gene (GLUT1). (K) Representation of ^13^C incorporation into glutamate omitting, for simplicity, generation of isotopomers from unlabeled pyruvate/acetyl-CoA, and respective representative multiplets observed in ^13^C NMR spectra measured in extracts after metabolizing [1-^13^C]glucose for 24 hours. (L) Glutamate (Glu) multiplet fractions, and fractional enrichment (FE) of lactate C3 of (Lac) and alanine (Ala). (M) Model used in the TCAcalc analysis and relative fluxes and lactate labeling estimated by fitting glutamate isotopomers, and Ala C3. Abbreviations: CS, citrate synthase; PDH, pyruvate dehydrogenase; Y, flux of anaplerotic substrates through pyruvate carboxylase (Y_PC_) or succinyl-CoA (Y_S_). Data is shown as mean±SD of 3-12 independent experiments, represented by the individual symbols. *P<0.05, **P<0.01, ***P<0.001 depict differences in comparisons following significant effects in ANOVA.

Then, we analyzed ECAR as a surrogate of glycolysis (Figure 2G-I). When compared to vehicle, both palmitate and LPS treatments increased basal glycolysis and the total glycolytic capacity, but only LPS increased the glycolytic reserve significantly (Figure 2I). Non-glycolytic medium acidification was similar across treatments. We further measured the expression of *Slc2a1*, the gene encoding the glucose carrier GLUT1, which tended to be increased by palmitate (adjusted P=0.065) and LPS (adjusted P=0.097) treatments compared to vehicle (ANOVA F(2,21)=3.54, P=0.048; Figure 1J).

Finally, we performed ^13^C metabolic tracing to infer on the rearrangement of fluxes through anaplerotic pathways. After treatment, cells were allowed to metabolize [1-^13^C]glucose for 24 hours, and metabolite extracts were analyzed by ^13^C NMR spectroscopy to determine labeling of glutamate carbons (Figure 2K). Glutamate (Glu) multiplet fractions, were mainly affected by LPS when compared to vehicle, namely Glu C3 (Figure 2L). However, a tendency for reduced labeling of multiplets in Glu C2 is apparent for palmitate *versus* vehicle. A metabolic flux analysis using TCAcalc estimated that, relative to the citrate synthase flux, pyruvate carboxylation was higher in palmitate-treated than vehicle, and blunted in LPS treated cells (Figure 2M). LPS-treated cells had, instead, faster rate of anaplerosis feeding succinyl-CoA than the remaining groups. Lactate labeling was a fitted parameter in the model, and the result correlated well with that measured experimentally (Pearson r=0.9998, P=0.014).

In sum, the increased proliferation induced by palmitate is accompanied by a rearrangement of energy metabolism fluxes, namely increased glycolysis, reduced mitochondrial respiration, and increased pyruvate carboxylation that can support *de novo* oxaloacetate synthesis for replenishment Krebs cycle intermediates used in biosynthetic pathways.

### Microglia communication via EVs

Palmitate-treated cells did not exacerbate cytokine production, we thus tested whether palmitate can modulate EVs as means of communication to other cells. Cells treated with either palmitate or LPS produced EVs of size similar to vehicle-treated cells, as determined by NTA (Figure 3A-B).

**Figure 3.**
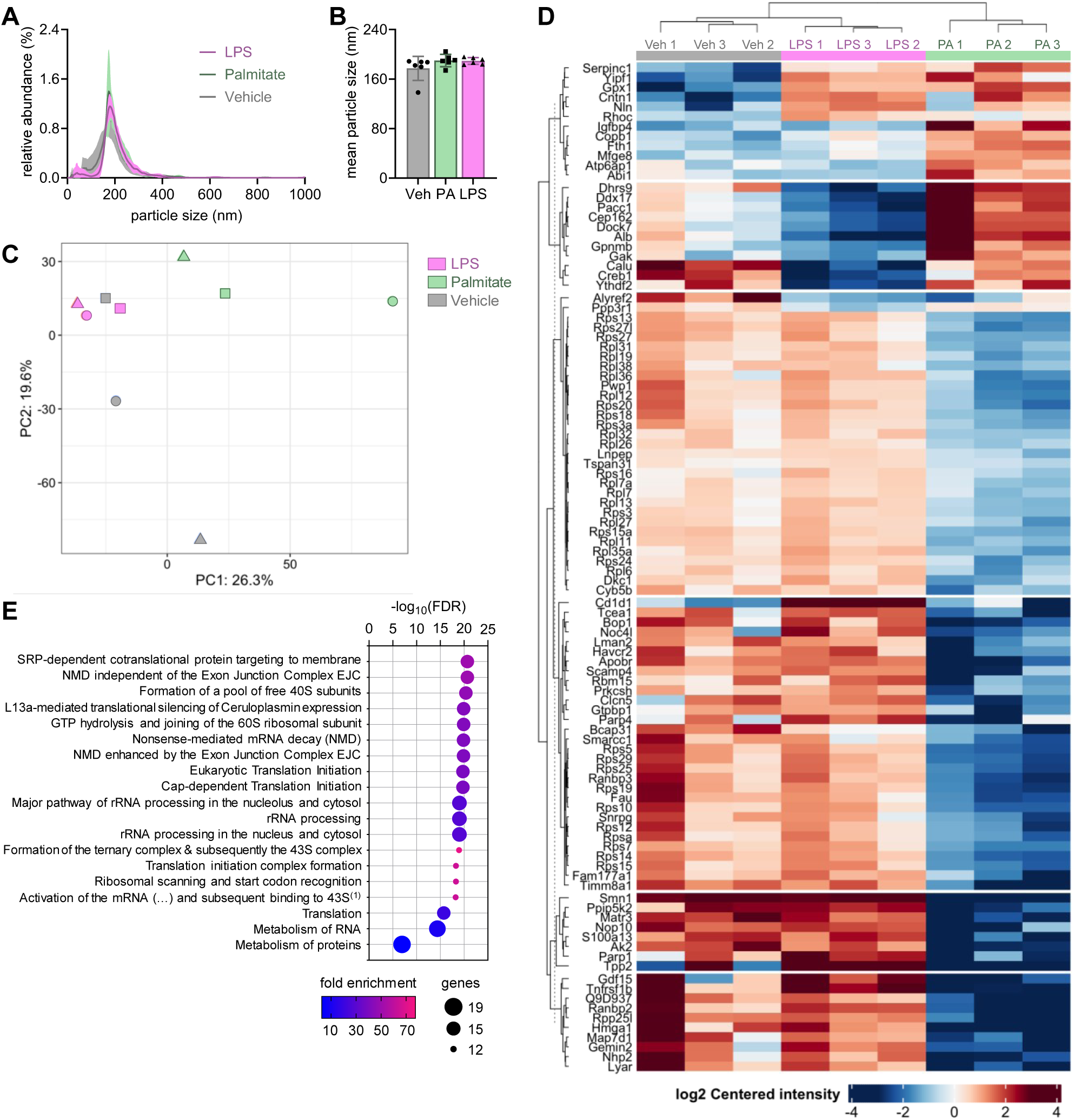
Palmitate alters the proteome of BV2-retreased EVs, including a reduction of proteins involved in RNA processing and protein synthesis. (A) Histograms of EV size distribution evaluated by Nanoparticle tracking analysis (NTA) of 6 independent EV isolations, and (B) estimated mean particle size for vehicle (Veh), palmitate (PA) and LPS. Data shown as mean±SD of n=6. (C) Score plots of a PCA of the 1000 most abundant proteins in EVs (n=3/group). Each symbol shape represents an independent experiment. (D) Heatmap of significant EV proteome differences between either of the 3 experimental groups, and (E) significant findings at FDR<0.01 from the gene ontology analysis of differentially expressed proteins between EVs from palmitate- and vehicle-treated BV2 cells. ^(1)^Full pathway name: “Activation of the mRNA upon binding of the cap-binding complex and eIFs and subsequent binding to 43S”.

We then investigated the proteome of microglia-derived EVs in response to each treatment. A false discovery rate (FDR) of <0.01 was set as a threshold protein identification, resulting in a total of 3739 proteins present across a total of 9 experiments (3 per treatment group). From these, only 1636 proteins were present in all analyzed samples. For subsequent analysis, only proteins that were present in at least 2 out of the 3 experiments of each group were kept.

Data was not found missing at random. In fact, EVs from palmitate-exposed cells seemed to lack of proteins that are present in the other groups. Moreover, missing proteins had lower average abundance in samples that did have a measurable signal, suggesting that these are most likely proteins that have very low abundance or are completely absent (true zeroes).

In a PCA using the 1000 most abundant proteins, PC1 allowed to separate palmitate from both LPS and vehicle treatments (Figure 3C), suggesting that 24 hours of exposure to palmitate triggers a unique proteomic signature of released EVs. A differential enrichment analysis revealed significant differences (adjusted P<0.05) in the abundance of 102 proteins in-between either of the 3 comparisons performed (figure 3D). Notably, palmitate-induced alterations on the EV proteome included a reduction in the abundance of several ribosomal proteins and proteins involved in protein metabolism. Indeed, a gene ontology analysis to the EV proteins that were differentially enriched between palmitate and vehicle treatments revealed pathways related to RNA processing, ribosome assembly, and translation processes (Figure 3E).

### EVs from palmitate-treated cells impact brain function

EVs were obtained from BV2 cells exposed to either palmitate or vehicle, and injected into the lateral ventricle mice, which allows spreading an infusate through the whole brain (confirmed in pilot experiments with trypan blue). A week after injection, we assessed exploratory behavior, memory performance, and depressive-like behavior (Figure 4A). The i.c.v. injection of EVs had no impact on body weight (Figure 4B), suggesting good recovery from surgery. In NOR and NOL, neither treatment group showed signs of object bias in the familiarization session (Figure 4C-D). In the recognition session, mice injected with EVs from vehicle-treated BV2 cells showed preserved memory performance as assessed in the NOR (P=0.029 *versus* random exploration) and NLR (P=0.010 *versus* random exploration). In turn, mice receiving EVs from BV2 cells exposed to palmitate did not show increased exploration of either the novel object (P=0.710 *versus* random exploration; Figure 4C) or novel location (P=0.117 *versus* random exploration; Figure 4C). When comparing memory performance between the treatment groups, significantly lower memory performance between palmitate and vehicle was observed in the NOR task (Figure 4C).

**Figure 4.**
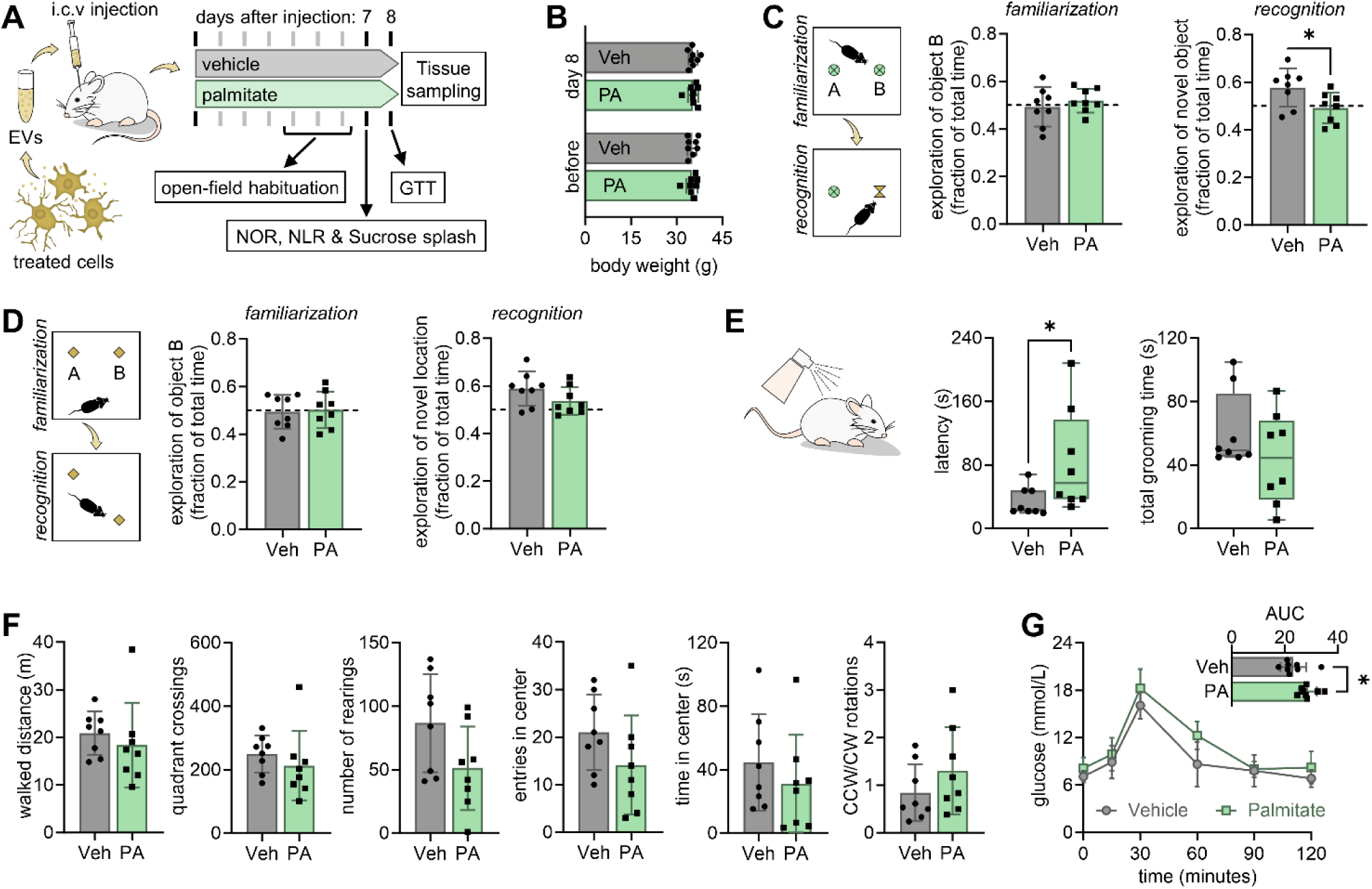
Intracerebroventricular (i.c.v) injection of microglia-derived EVs following palmitate exposure affects cognition and depressive-like behavior, and alters glucose metabolism in mice. (A) Mice were injected in the lateral ventricle with EVs (500 ng of protein) collected from BV2 cells after exposure to either palmitate (PA, 200 µmol/L) or vehicle (Veh). (B) Body weight of mice before and 8 days after surgery for EV administration. Impaired memory was observed novel object recognition (NOR) test (C), and a similar trend was observed in the novel location recognition (NLR) test (D). (E) Depression-related behavior was assessed by grooming behavior in the sucrose splash test. (F) locomotor activity and exploratory behavior assessed in the last habituation day to the open-field arena. (G) Glucose clearance in the GTT in the 8^th^ day following EV injection. The inset is the area under the curve (AUC) of the glucose excursion for each mouse. Symbols representing each mouse (n=8) are overlaid on bar graphs showing mean±SD or box plots showing interquartile ranges. *P<0.05 from either Student’s t-test or with Mann Whitney test.

The sucrose splash test is used to infer on self-care behavior. Mice injected i.c.v. with EVs from palmitate-exposed microglia showed significantly larger latency to start grooming, when compared to vehicle (P=0.037; Figure 4E). Such reduced self-care behavior is characteristic of depression, although there was no significant change on the total duration of grooming between treatments.

A number of parameters of locomotion and exploratory behavior were measured in the empty open field arena, showing no significant differences between the treatments (Figure 4F). Notably, the ratio of counterclockwise-to-clockwise rotations was also similar between groups and did not show signs of lateralized motor impairment (Figure 4F).

### EVs impact glucose homeostasis

EVs injected i.c.v. may impact the hypothalamus, or even reach the blood stream and act on peripheral organs (Banks *et al*., 2020), thus affecting metabolism and glucose homeostasis. Therefore, we performed a GTT after behavior assessments. While baseline glucose was similar between treatment groups (P=0.083), mice receiving EVs from palmitate-exposed BV2 cells showed a larger glucose excursion than the vehicle group, as typified by increased GTT area-under the curve (+24%, P=0.018; Figure 4G).

### EVs can impact microglia in the mouse brain

EVs are likely to afford exchange of cellular material in-between microglia, and from microglia to other cells. We then tested whether the injected EVs induced alterations in the morphology of microglia and astrocytes in the mouse hypothalamus (*arcuate nucleus*, ARC), and in the hippocampal *cornus ammonis* (CA1 and CA3) and *dentate gyrus* (DG) (Figure 5A). We determined the area covered by immunoreactivity against Iba1 and GFAP, which are microglia and astrocyte markers, respectively. Compared to vehicle, mice treated with EVs from palmitate-exposed BV2 cells showed significantly larger Iba1 coverage across all regions analyzed (ANOVA: region x treatment F(3,36)=0.064, P=0.979; region F(3,36)=1.081, P=0.369; treatment F(1,12)=14.250, P=0.003; Figure 5B). In contrast, GFAP area was similar in the 2 groups, suggesting that despite microglia expansion, the EVs from palmitate-treated BV2 microglia did not cause overt gliosis *in vivo* (Figure 5C). Next, we set to explore the morphology of Iba1^+^ cells, that is microglia. Microglia from the brain of mice in palmitate and vehicle groups showed similar number of processes sprouting from the cell soma (Figure 5D). In turn, compared to vehicle, mice injected i.c.v. with EVs from palmitate-exposed cells showed more branching points in the microglia processes (ANOVA: region x treatment F(3,36)=0.142, P=0.934; region F(3,36)=0.546, P=0.654; treatment F(1,12)=5.141, P=0.043; Figure 5E). Total length and maximum length of the cell processes were similar between groups (Figure 5F-G).

**Figure 5.**
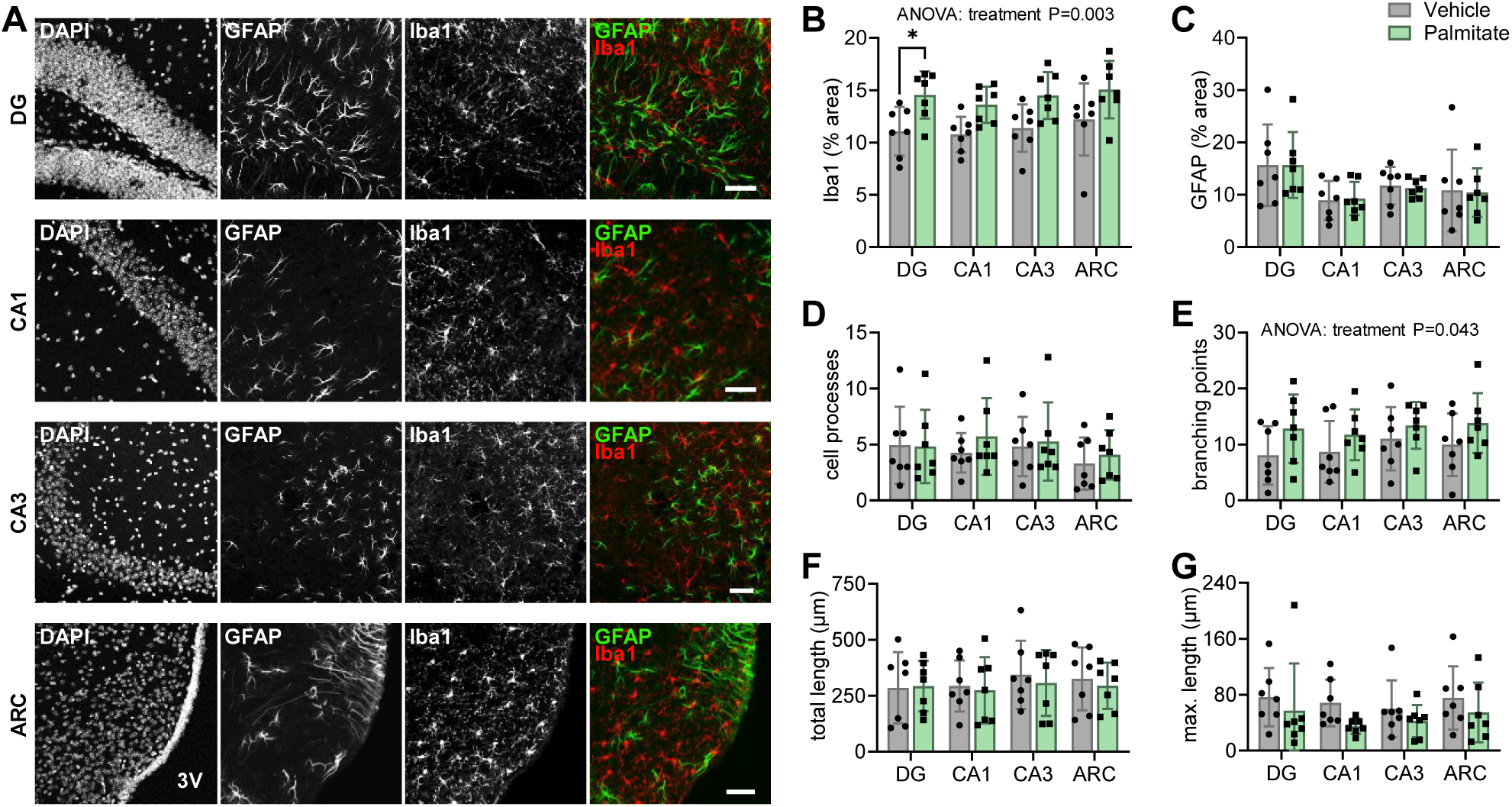
EVs from palmitate-challenged microglia *in vitro* activate microglia in the hippocampus and hypothalamus of mice *in vivo*. (A) Representative micrographs showing DAPI signal from nuclei, GFAP immunoreactivity in red, and Iba1 immunoreactivity in red in the *dentate gyrus* (DG) and *cornus ammonis* (CA1/CA3) areas of the hippocampus, and *arcuate nucleus* (ARC) of the hypothalamus at 8 days after intraventricular EVs injection. Scale bars over the micrographs indicate 50 µm; 3V=third ventricle. (B-C) Fraction of area occupied by iba1 and GFAP immunoreactivity in DG, CA1, CA3 and ARC. (D) Mean number of processes, (E) number of branching points, (F) total cell process length, and (G) maximum process length of microglia, as determined from skeleton analysis of 3-4 iba1^+^ cells per mouse. Data is shown as mean±SD of 7 independent experiments, represented by the individual symbols. *P<0.05 depicts differences in comparisons following significant effects in ANOVA.

These findings indicate that EVs shed by palmitate-exposed microglia induce microglia branching *in vivo*, despite no overt gliosis after one week of direct administration to the brain.

## Discussion

The present results suggest that microglia respond to palmitate exposure with increased proliferation, and with a metabolic network rearrangement that favors energy production from glycolysis rather than oxidative metabolism, despite stimulated mitochondria biogenesis. In addition, while palmitate did not induce increased cytokine expression, it modified the protein cargo of released EVs which, alone, can contribute to the development of memory impairment, depression-like behavior, and glucose intolerance.

### Rearrangement of energy metabolism after palmitate exposure

Microglia are metabolically flexible cells that can utilize a variety of substrates, and their activation upon injury or infection is an energetically costly process (Bernier *et al*., 2020; reviewed in Aldana, 2019). When activated by inflammatory stimuli, microglial cells exacerbate glucose uptake and glycolysis (Ghosh *et al*., 2018; Gimeno-Bayón *et al*., 2014). A similar metabolic shift occurs in activated macrophages (Galván-Peña & O’Neill, 2014). Indeed, we have observed this switch from oxidative metabolism to anaerobic glycolysis in BV2 microglia activated by either palmitate or LPS. In contrast, Chausse *et al*. have reported that BV2 activation by palmitate is sustained by oxidative metabolism, while LPS activation is mostly dependent on glycolysis (Chausse *et al*., 2019). It should be noted, however, that Chausse *et al*. have cultured their cells in the presence of 25 mmol/L of glucose, which is believed to represent hyperglycemic conditions (*e.g.*, see Duarte *et al*., 2009 for brain glucose levels observed in the brain of humans and rodents). Primary microglia and BV2 microglial cells express several glucose carriers, with GLUT1 being at least one order of magnitude more abundant than the others, and the one that is upregulated upon microglia activation (Wang *et al*., 2019). Our results also support GLUT1 upregulation in both LPS and palmitate exposure.

Activation of BV2 cells and primary microglia by LPS is known to induce mitochondrial fragmentation (Park *et al*., 2013; Nair *et al*., 2018). Enhanced expression of genes coding for proteins involved in mitochondria fragmentation was not observed in our short (3 hours) LPS exposure or upon palmitate treatment. Namely, mRNA levels of mitochondrial fission markers dynamin-related protein 1(Drp1), fission protein 1 (Fis1), and mitochondrial fission factor (Mff) were similar across experimental groups. Instead, we observed a palmitate-induced expression of PGC-1α, which is a key regulator of mitochondrial biogenesis. Indeed, palmitate also induced an increase in mitochondria content, and in the density of proteins belonging to the mitochondrial complexes of the electron transport chain. Together, our findings suggest that although palmitate increases mitochondria biogenesis, it reduces the efficiency of respiration and exacerbates glycolysis. Using ^13^C tracing experiments, we further found that palmitate further increases the rate of pyruvate carboxylation, which is needed for *de novo* oxaloacetate synthesis (Rae *et al*., 2024), and supports the drainage of Krebs cycle intermediates used in biosynthetic pathways during increased cellular proliferation.

### Lack of palmitate-induced cytokine overexpression

Obesity and saturated fat exposure is known to stimulate NF-κB transcriptional activity and the secretion of pro-inflammatory mediators in peripheral organs, such as liver, white adipose tissue, and leukocytes (Saltiel & Olefsky, 2017). In the brain, the neuroinflammatory process in DIO remains unclear, and somehow controversial. That might be because microglia reactivity is modulated by specific neurochemical signals within the cell’s microenvironment, and due to transient release of inflammatory mediators. Indeed, there have been reports of both differential microgliosis profiles across distinct brain areas (Brandi *et al*., 2022; Mrdjen *et al*., 2023), and biphasic gliosis with transient cytokine release in chronic noxious stimuli, such as that in DIO (Thaler *et al*., 2012; de Paula *et al*., 2021).

We have conducted this study under the assumption that palmitate is the saturated fatty acid in the context of obesity driving neuroinflammation (see Melo *et al*., 2020). In fact, palmitate induces much stronger expression of inflammatory factors in primary astrocytes than saturated fatty acids such as laurate or stearate (Gupta *et al*., 2012). In addition, 200 µmol/L of palmitate but not stearate impairs mitochondrial function in the C6 astrocyte cell line (Schmitt *et al*., 2024). In neuronal cell models, both palmitate and stearate are able to cause cell death (Ulloth *et al*., 2003). In our hands, while the exposure of BV2 cells to LPS resulted in an enormous expression of pro-inflammatory cytokines, palmitate was devoid of such effect. Accordingly, Chausse *et al*. also failed to find increased inflammatory mediators in palmitate- or oleate-exposed BV2 cells (Chausse *et al*., 2019). Others have suggested that palmitate can even activate anti-inflammatory pathways and reduce TNF-α expression in BV2 microglia (Tracy *et al*., 2013; Kim *et al*., 2018). Despite not significantly different, our data also suggests a transient reduction of IL-6 expression at the beginning of palmitate exposure, when compared to vehicle-treated BV2 cells (see Figure 1F).

### EVs as neuroinflammatory mediators

DIO leads to depression-like behavior and hippocampal-dependent memory impairment in rodents, although not all diet intervention studies have found pronounced increases in neuroinflammatory markers (Pistell *et al*., 2010; Kaczmarczyk *et al*., 2013; Mansur *et al*., 2015; Lizarbe, Soares *et al*., 2019; Garcia-Serrano, Mohr *et al*., 2022; Garcia-Serrano, Vieira *et al*., 2022; Skoug *et al*., 2024; de Paula *et al*., 2021). Microglial inhibition protocols have supported the notion that microgliosis is an important contributor to neuronal dysfunction in the *arcuate nucleus* of the hypothalamus upon DIO (Valdearcos *et al*., 2014), and can both prevent peripheral inflammation and reduce the extent of metbolic syndrome development (André *et al*., 2017).

However, the exacerbated cytokine production appears to be transient in both the hippocampus and hypothalamus during DIO (Thaler et al., 2012, de Paula *et al*., 2021). In later stages of obesity development, in the absence of cytokine overproduction, EVs could contribute with molecular messages from microglia to other cells. EVs isolated from palmitate- or LPS-treated cells had unaltered size but displayed significant proteome changes. Interestingly, our pattern recognition analysis demonstrated larger proteome differences between palmitate and vehicle than between LPS and vehicle groups.

Most significant differentially expressed proteins in palmitate were related to mRNA processing and translation into protein, including the downregulation of several ribosome subunits (Rps5, Rps7, Rps13, Rps14, Rps15a, Rps29, Rpl11, Rpl35a). Ribosome subunits in neurons exhibit a dynamic assembly that is locally regulated in dendrites and synapses (Shigeoka *et al*., 2019; Fusco *et al*., 2021). Moreover, RNA stress granules and smaller dendritic trees were previously observed when ribosomal proteins were depleted from neurons with already established dendrites (Slomnicki *et al*., 2016). Thus, it is likely that microglia EVs deliver components of the ribosomal machinery to support protein synthesis in neuronal processes, and we speculate that the reduction in such components within microglial EVs after palmitate exposure can result in alterations of synaptic plasticity and neuronal connectivity. Decreased expression of ribosomal proteins have been reported in metabolically healthy obese individuals (Gaye *et al*., 2018), including Rps29 and Rpl35a, also found to be downregulated in the present study.

Previous reports demonstrated that EVs from microglia activated by ATP, IFN-γ or LPS can impact neurons and astrocytes (Drago *et al*., 2017; Tsutsumi *et al*., 2019). In our experimental setup, when EVs were given i.c.v., mice receiving EVs from palmitate-treated microglia developed memory impairment, as assessed by the lower capacity to recognize the novelty in NOR and NLR tests. An increase in latency to grooming in the sucrose splash test further indicated a depression-like behavior. Moreover, mice injected with EVs derived from palmitate-exposed microglia developed glucose intolerance, when compared to the vehicle group, suggesting alterations in central regulation of metabolism. In DIO, impaired glucose homeostasis occurs before insulin resistance is installed (Soares *et al*., 2018; Garcia-Serrano, Mohr *et al*., 2022), which is in line with direct actions of EVs on hypothalamic glucose-sensing neurons. However, one must keep in mind that smaller EVs, such as exosomes, are known to cross the blood-brain barrier (Winston *et al*., 2019; Zhang *et al*., 2019; Wiklander *et al*., 2019; Serpe *et al*., 2021), and could act on peripheral organs as well.

Finally, we investigated whether EVs alone could be mediators of neuroinflammation. Thus, we evaluated gliosis in the mouse brain following EV administration, with focus on the hippocampus that controls learning and memory, and the hypothalamus that is the main central glucose sensing area. Notably, we have observed some degree of microgliosis but not astrogliosis when injecting i.c.v. the EVs from palmitate-exposed microglia, relative to EVs from vehicle experiments. The absence of astrogliosis is somehow surprising since microglia-derived EVs after ATP stimulation were found to modulate astrocytes (Drago *et al*., 2017).

### Limitations

We have characterized energy metabolism of BV2 cells, but have not performed in-depth investigations into the mechanisms that drive mitochondrial alterations, which could include intracellular accumulation of lipids, and the impairment in insulin signaling. Moreover, although this study puts forward novel ideas for further testing, it is important to note that we have studied the BV2 microglial cell line and not primary microglia. The cell line allowed us to have access to larger amounts of material (namely EVs) without collecting tissue from large numbers of animals, but our findings might not fully represent microglia responses *in vivo*. In addition, we have studied protein content of EVs, but not other types of cargo, such as small metabolites, lipids, and nucleic acids. In fact, it is well known that EVs carry microRNAs that control the expression of a variety of proteins, namely proteins involved in synaptic pruning, energy metabolism, immune response, and translation/transcription (Prada *et al*., 2018; Yang *et al*., 2018; Karvinen *et al*., 2023). Whether these microRNAs are present and modified after palmitate exposure was not tested, but further studies on this topic are warranted.

## Conclusion

In this study, using BV2 microglia, we demonstrate that palmitate exposure does not drive expression of cytokines that is typical of microglia activation. Instead, palmitate led to the release of EVs with modified cargo that, alone, is sufficient to induce some degree of microgliosis in the mouse, and to impact brain function. The EV-mediated palmitate dysfunction is thus a plausible mechanism by which microglia participate in brain dysfunction during DIO.

## Author Contributions

JMND designed the study. GCdP, RB, RF-C, AB and BA carried out experiments. IL, TD and JMND provided resources and supervised experimental work. GCdP and JMND wrote the manuscript. All authors revised the manuscript.

## Conflict of interest disclosure

The authors declare no competing interests in relation to this work.

## Data availability

The datasets from the current study are available from the corresponding author on reasonable request.

## Ethics approval statement

Experiments were approved by the Malmö-Lund Committee for Animal Experiment Ethics (#5123-21).

## Acknowledgements

The authors thank Charlotte Welinder for the mass spectrometry analysis, Daria Rago for help with the proteomic analysis, and Patricia F. Nunes (The Francis Crick Institute) for discussions on palmitate preparation and cell treatments. We acknowledge the Strategic Research Area MultiPark (Multidisciplinary Research on Parkinson’s Disease) for access to mouse behavior labs, the Lund University Bioimaging Centre for access to microscopy resources, and the BioMS for access to proteomics facility.

This work was supported by the Swedish foundation for International Cooperation in Research and Higher education (#BR2019-8508), Swedish Research Council (#2019-01130), Diabetesfonden (#Dia2019-440, #Dia2021-637), Direktör Albert Påhlssons Foundation, and Royal Physiographic Society of Lund. JMND acknowledges support from The Knut and Alice Wallenberg foundation, infrastructure funding of Lund University (Dnr STYR 2019/318) and Lund University Faculty of Medicine (Dnr STYR 2021/2984), and Lund University Diabetes Centre, which is funded by the Swedish Research Council (Strategic Research Area EXODIAB; grant no.: 2009-1039) and the Swedish Foundation for Strategic Research (grant no.: IRC15-0067). RB and IL are funded by the Parkinson Research Foundation.

## Notes

### Competing Interest Statement

The authors have declared no competing interest.

## References

1. Aldana, B.I. (2019). Microglia-Specific Metabolic Changes in Neurodegeneration. J Mol Biol. 431(9):1830–1842. doi: 10.1016/j.jmb.2019.03.006.

2. Alger JR, Minhajuddin A, Dean Sherry A, Malloy CR. Analysis of steady-state carbon tracer experiments using akaike information criteria. Metabolomics. 2021 Jun 19;17(7):61. doi: 10.1007/s11306-021-01807-1. PMID: 34148138.

3. André, C., Guzman-Quevedo, O., Rey, C., Rémus-Borel, J., Clark, S., Castellanos-Jankiewicz, A., Ladeveze, E., Leste-Lasserre, T., Nadjar, A., Abrous, D.N., Laye, S., Cota, D. (2017). Inhibiting Microglia Expansion Prevents Diet-Induced Hypothalamic and Peripheral Inflammation. Diabetes. 66(4):908–919. doi: 10.2337/db16-0586.

4. Arganda-Carreras, I., Fernández-González, R., Muñoz-Barrutia, A., & Ortiz-De-Solorzano, C. (2010). 3D reconstruction of histological sections: Application to mammary gland tissue. Microscopy Research and Technique, 73(11), 1019–1029. doi:10.1002/jemt.20829

5. Attuquayefio TN, Stevenson RJ, Oaten MJ, Francis HM. (2017). A four-day Western-style dietary intervention causes reductions in hippocampaldependent learning and memory and interoceptive sensitivity. PLoS One 12:e0172645. doi: 10.1371/journal.pone.0172645.

6. Banks WA, Sharma P, Bullock KM, Hansen KM, Ludwig N, Whiteside TL. Transport of Extracellular Vesicles across the Blood-Brain Barrier: Brain Pharmacokinetics and Effects of Inflammation. Int J Mol Sci. 2020 Jun 21;21(12):4407. doi: 10.3390/ijms21124407. PMID: 32575812; PMCID: PMC7352415.

7. Baufeld C, Osterloh A, Prokop S, Miller KR, Heppner FL. High-fat diet-induced brain region-specific phenotypic spectrum of CNS resident microglia. Acta Neuropathol. 2016 Sep;132(3):361–75. doi: 10.1007/s00401-016-1595-4. Epub 2016 Jul 8. PMID: 27393312; PMCID: PMC4992033.

8. Bernhart, E., Kollroser, M., Rechberger, G., Reicher, H., Heinemann, A., Schratl, P., . . . Sattler, W. (2010). Lysophosphatidic acid receptor activation affects the C13NJ microglia cell line proteome leading to alterations in glycolysis, motility, and cytoskeletal architecture. Proteomics, 10, 141–158. doi: 10.1002/pmic.200900195.

9. Bernier LP, York EM, Kamyabi A, Choi HB, Weilinger NL, MacVicar BA. Microglial metabolic flexibility supports immune surveillance of the brain parenchyma. Nat Commun. 2020 Mar 25;11(1):1559. doi: 10.1038/s41467-020-15267-z. PMID: 32214088; PMCID: PMC7096448.

10. Brandi E, Torres-Garcia L, Svanbergsson A, Haikal C, Liu D, Li W, Li JY. Brain region-specific microglial and astrocytic activation in response to systemic lipopolysaccharides exposure. Front Aging Neurosci. 2022 Aug 26;14:910988. doi: 10.3389/fnagi.2022.910988. PMID: 36092814; PMCID: PMC9459169.

11. Caruso Bavisotto C, Scalia F, Marino GA, Carlisi D, Bucchieri F, Conway de Macario E. (2019). Extracellular Vesicle-Mediated Cell⁻Cell Communication in the Nervous System: Focus on Neurological Diseases. Int J Mol Sci. 20(2):434. doi: 10.3390/ijms20020434.

12. Cavaliere, G., Trinchese, G., Penna, E., Cimmino, F., Pirozzi, C., Lama, A., … Mollica, M.P. (2019). High-Fat Diet Induces Neuroinflammation and Mitochondrial Impairment in Mice Cerebral Cortex and Synaptic Fraction. Front Cell Neurosci. 13:509. doi: 10.3389/fncel.2019.00509.

13. Chausse, B., Kakimoto, P.A., Caldeira-da-Silva, C.C., Chaves-Filho, A.B., Yoshinaga, M.Y., da Silva, R.P., Miyamoto, S., Kowaltowski, A.J. (2019). Distinct metabolic patterns during microglial remodeling by oleate and palmitate. Biosci Rep. 39(4):BSR20190072. doi: 10.1042/BSR20190072.

14. de Paula GC, Brunetta HS, Engel DF, Gaspar JM, Velloso LA, Engblom D, de Oliveira J, de Bem AF. Hippocampal Function Is Impaired by a Short-Term High-Fat Diet in Mice: Increased Blood-Brain Barrier Permeability and Neuroinflammation as Triggering Events. Front Neurosci. 2021 Nov 4;15:734158. doi: 10.3389/fnins.2021.734158. PMID: 34803583; PMCID: PMC8600238.

15. Drago F, Lombardi M, Prada I, Gabrielli M, Joshi P, Cojoc D, Franck J, Fournier I, Vizioli J, Verderio C. (2017). ATP Modifies the Proteome of Extracellular Vesicles Released by Microglia and Influences Their Action on Astrocytes. Front Pharmacol. 8:910. doi: 10.3389/fphar.2017.00910.

16. Duarte JMN, Morgenthaler FD, Lei H, Poitry-Yamate C, Gruetter R. Steady-state brain glucose transport kinetics re-evaluated with a four-state conformational model. Front Neuroenergetics. 2009 Oct 12;1:6. doi: 10.3389/neuro.14.006.2009. PMID: 20027232; PMCID: PMC2795468.

17. Duarte JMN, Cunha RA, Carvalho RA. Different metabolism of glutamatergic and GABAergic compartments in superfused hippocampal slices characterized by nuclear magnetic resonance spectroscopy. Neuroscience. 2007 Feb 23;144(4):1305–13. doi: 10.1016/j.neuroscience.2006.11.027. Epub 2006 Dec 29. PMID: 17197104.

18. Duarte JMN. Loss of brain energy metabolism control as a driver for memory impairment upon insulin resistance. Biochem Soc Trans. 2023 Feb 27;51(1):287–301. doi: 10.1042/BST20220789. PMID: 36606696.

19. Egnatchik RA, Leamy AK, Noguchi Y, Shiota M, Young JD. Palmitate-induced activation of mitochondrial metabolism promotes oxidative stress and apoptosis in H4IIEC3 rat hepatocytes. Metabolism. 2014 Feb;63(2):283–95. doi: 10.1016/j.metabol.2013.10.009. Epub 2013 Oct 24. PMID: 24286856; PMCID: PMC3946971.

20. Fusco CM, Desch K, Dörrbaum AR, Wang M, Staab A, Chan ICW, Vail E, Villeri V, Langer JD, Schuman EM. Neuronal ribosomes exhibit dynamic and context-dependent exchange of ribosomal proteins. Nat Commun. 2021 Oct 21;12(1):6127. doi: 10.1038/s41467-021-26365-x. PMID: 34675203; PMCID: PMC8531293.

21. Galván-Peña, S., O’Neill, L.A. (2014). Metabolic reprograming in macrophage polarization. Front Immunol. 5:420. doi: 10.3389/fimmu.2014.00420.

22. Garcia-Serrano AM, Duarte JMN. Brain Metabolism Alterations in Type 2 Diabetes: What Did We Learn From Diet-Induced Diabetes Models? Front Neurosci. 2020 Mar 20;14:229. doi: 10.3389/fnins.2020.00229. PMID: 32265637; PMCID: PMC7101159.

23. Garcia-Serrano AM, Mohr AA, Philippe J, Skoug C, Spégel P, Duarte JMN. Cognitive Impairment and Metabolite Profile Alterations in the Hippocampus and Cortex of Male and Female Mice Exposed to a Fat and Sugar-Rich Diet are Normalized by Diet Reversal. Aging Dis. 2022 Feb 1;13(1):267–283. doi: 10.14336/AD.2021.0720. PMID: 35111373; PMCID: PMC8782561.

24. Garcia-Serrano AM, Vieira JPP, Fleischhart V, Duarte JMN. Taurine and N-acetylcysteine treatments prevent memory impairment and metabolite profile alterations in the hippocampus of high-fat diet-fed female mice. Nutr Neurosci. 2023 Nov;26(11):1090–1102. doi: 10.1080/1028415X.2022.2131062. Epub 2022 Oct 12. PMID: 36222315.

25. Gaye A, Doumatey AP, Davis SK, Rotimi CN, Gibbons GH. (2018). Whole-genome transcriptomic insights into protective molecular mechanisms in metabolically healthy obese African Americans. NPJ Genom Med. 3:4. doi: 10.1038/s41525-018-0043-x.

26. Ghosh, S., Castillo, E., Frias, E.S., Swanson, R.A. (2018). Bioenergetic regulation of microglia. Glia. 66(6):1200–1212. doi: 10.1002/glia.23271.

27. Gimeno-Bayón, J., López-López, A., Rodríguez, M.J., Mahy, N. (2014). Glucose pathways adaptation supports acquisition of activated microglia phenotype. J Neurosci Res. 92 (6):723–31. doi: 10.1002/jnr.23356.

28. Gupta S, Knight AG, Gupta S, Keller JN, Bruce-Keller AJ. Saturated long-chain fatty acids activate inflammatory signaling in astrocytes. J Neurochem. 2012 Mar;120(6):1060–71. doi: 10.1111/j.1471-4159.2012.07660.x. Epub 2012 Feb 6. PMID: 22248073; PMCID: PMC3296820.

29. Halling JF, Pilegaard H. PGC-1α-mediated regulation of mitochondrial function and physiological implications. Appl Physiol Nutr Metab. 2020 Sep;45(9):927–936. doi: 10.1139/apnm-2020-0005. Epub 2020 Jun 9. PMID: 32516539.

30. Hartmann, H., Pauli, L.K., Janssen, L.K., Huhn, S., Ceglarek, U., Horstmann, A. (2020). Preliminary evidence for an association between intake of high-fat high-sugar diet, variations in peripheral dopamine precursor availability and dopamine-dependent cognition in humans. J Neuroendocrinol. (12):e12917. doi: 10.1111/jne.12917.

31. Hoscheidt, S., Sanderlin, A.H., Baker, L.D., Jung, Y., Lockhart, S., Kellar, D., Whitlow, C.T., Hanson, A.J., Friedman, S., Register, T., Leverenz, J.B., Craft, S. (2022). Mediterranean and Western diet effects on Alzheimer’s disease biomarkers, cerebral perfusion, and cognition in mid-life: A randomized trial. Alzheimers Dement. (3):457–468. doi: 10.1002/alz.12421.

32. Hu S, Hu Y, Yan W. Extracellular vesicle-mediated interorgan communication in metabolic diseases. Trends Endocrinol Metab. 2023 Sep;34(9):571–582. doi: 10.1016/j.tem.2023.06.002. Epub 2023 Jun 30. PMID: 37394346.

33. Kaczmarczyk, M.M., Machaj, A.S., Chiu, G.S., Lawson, M.A., Gainey, S.J., York, J.M., et al. (2013). Methylphenidate prevents high-fat diet (HFD)-induced learning/memory impairment in juvenile mice. Psychoneuroendocrinology. 38, 1553–1564. doi: 10.1016/j.psyneuen.2013.01.004.

34. Karvinen S, Korhonen TM, Sievänen T, Karppinen JE, Juppi HK, Jakoaho V, Kujala UM, Laukkanen JA, Lehti M, Laakkonen EK. (2023). Extracellular vesicles and high-density lipoproteins: Exercise and oestrogen-responsive small RNA carriers. J Extracell Vesicles. 12(2):e12308. doi: 10.1002/jev2.12308.

35. Kim, S.M., McIlwraith, E.K., Chalmers, J.A., Belsham, D.D. (2018). Palmitate Induces an Anti-Inflammatory Response in Immortalized Microglial BV-2 and IMG Cell Lines that Decreases TNFα Levels in mHypoE-46 Hypothalamic Neurons in Co-Culture. Neuroendocrinology. 107(4):387–399. doi: 10.1159/000494759.

36. Lizarbe B, Cherix A, Duarte JMN, Cardinaux JR, Gruetter R. High-fat diet consumption alters energy metabolism in the mouse hypothalamus. Int J Obes (Lond). 2019 Jun;43(6):1295–1304. doi: 10.1038/s41366-018-0224-9. Epub 2018 Oct 9. PMID: 30301962.

37. Lizarbe B, Soares AF, Larsson S, Duarte JMN. Neurochemical Modifications in the Hippocampus, Cortex and Hypothalamus of Mice Exposed to Long-Term High-Fat Diet. Front Neurosci. 2019 Jan 8;12:985. doi: 10.3389/fnins.2018.00985. PMID: 30670942; PMCID: PMC6331468.

38. Mansur, R.B., Brietzke, E., McIntyre, R.S. (2015). Is there a “metabolic-mood syndrome”? A review of the relationship between obesity and mood disorders. Neurosci. Biobehav. Rev. 52, 89–104. doi: 10.1016/j.neubiorev.2014.12.017.

39. Melo, H.M., Seixas da Silva, G.D.S., Sant’Ana, M.R., Teixeira, C.V.L., Clarke, J.R., Miya Coreixas, V.S., et al. (2020). Palmitate Is Increased in the Cerebrospinal Fluid of Humans with Obesity and Induces Memory Impairment in Mice via Pro-inflammatory TNF-α. Cell Rep. 30(7):2180–2194.e8. doi: 10.1016/j.celrep.2020.01.072.

40. Mrdjen D, Amouzgar M, Cannon B, Liu C, Spence A, McCaffrey E, Bharadwaj A, Tebaykin D, Bukhari S, Hartmann FJ, Kagel A, Vijayaragavan K, Oliveria JP, Yakabi K, Serrano GE, Corrada MM, Kawas CH, Camacho C, Bosse M, Tibshirani R, Beach TG, Angelo M, Montine T, Bendall SC. Spatial proteomics reveals human microglial states shaped by anatomy and neuropathology. Res Sq [Preprint]. 2023 Jun 2:rs.3.rs-2987263. doi: 10.21203/rs.3.rs-2987263/v1. PMID: 37398389; PMCID: PMC10312937.

41. Nair S, Sobotka KS, Joshi P, Gressens P, Fleiss B, Thornton C, Mallard C, Hagberg H. Lipopolysaccharide-induced alteration of mitochondrial morphology induces a metabolic shift in microglia modulating the inflammatory response in vitro and in vivo. Glia. 2019 Jun;67(6):1047–1061. doi: 10.1002/glia.23587. Epub 2019 Jan 13. PMID: 30637805.

42. Park J, Choi H, Min JS, Park SJ, Kim JH, Park HJ, Kim B, Chae JI, Yim M, Lee DS. Mitochondrial dynamics modulate the expression of pro-inflammatory mediators in microglial cells. J Neurochem. 2013 Oct;127(2):221–32. doi: 10.1111/jnc.12361. Epub 2013 Jul 29. PMID: 23815397.

43. Park J., Choi H., Min J.-S., Park S.-J., Kim J.-H., Park H.J., Kim B., Chae J.-I., Yim M., Lee H.-J. (2013) Mitochondrial dynamics modulate the expression of pro-inflammatory mediators in microglial cells. J. Neurochem. 127:221–232. doi: 10.1111/jnc.12361.

44. Pascual M, Ibáñez F, Guerri C. (2021). Exosomes as mediators of neuron-glia communication in neuroinflammation. Neural Regen Res. 15(5):796–801. doi: 10.4103/1673-5374.268893.

45. Pistell, P.J., Morrison, C.D., Gupta, S., Knight, A.G., Keller, J.N., Ingram, D.K., Bruce-Keller, A.J. (2010). Cognitive impairment following high fat diet consumption is associated with brain inflammation. J Neuroimmunol. 219(1-2):25–32. doi: 10.1016/j.jneuroim.2009.11.010.

46. Prada I, Gabrielli M, Turola E, Iorio A, D’Arrigo G, Parolisi R, De Luca M, Pacifici M, Bastoni M, Lombardi M, Legname G, Cojoc D, Buffo A, Furlan R, Peruzzi F, Verderio C. (2018). Glia-to-neuron transfer of miRNAs via extracellular vesicles: a new mechanism underlying inflammation-induced synaptic alterations. Acta Neuropathol. 135(4):529–550. doi: 10.1007/s00401-017-1803-x.

47. Rae CD, Baur JA, Borges K, Dienel G, Díaz-García CM, Douglass SR, Drew K, Duarte JMN, Duran J, Kann O, Kristian T, Lee-Liu D, Lindquist BE, McNay EC, Robinson MB, Rothman DL, Rowlands BD, Ryan TA, Scafidi J, Scafidi S, Shuttleworth CW, Swanson RA, Uruk G, Vardjan N, Zorec R, McKenna MC. Brain energy metabolism: A roadmap for future research. J Neurochem. 2024 Jan 6. doi: 10.1111/jnc.16032. Epub ahead of print. PMID: 38183680.

48. Saltiel, A.R., Olefsky, J.M. (2017). Inflammatory mechanisms linking obesity and metabolic disease. J Clin Invest. 127(1):1–4. doi: 10.1172/JCI92035.

49. Serpe C, Monaco L, Relucenti M, Iovino L, Familiari P, Scavizzi F, Raspa M, Familiari G, Civiero L, D’Agnano I, Limatola C, Catalano M. (2021). Microglia-Derived Small Extracellular Vesicles Reduce Glioma Growth by Modifying Tumor Cell Metabolism and Enhancing Glutamate Clearance through miR-124. Cells 10(8):2066. doi: 10.3390/cells10082066.

50. Shigeoka T, Koppers M, Wong HH, Lin JQ, Cagnetta R, Dwivedy A, de Freitas Nascimento J, van Tartwijk FW, Ströhl F, Cioni JM, Schaeffer J, Carrington M, Kaminski CF, Jung H, Harris WA, Holt CE. On-Site Ribosome Remodeling by Locally Synthesized Ribosomal Proteins in Axons. Cell Rep. 2019 Dec 10;29(11):3605–3619.e10. doi: 10.1016/j.celrep.2019.11.025. PMID: 31825839; PMCID: PMC6915326.

51. Skoug C, Holm C, Duarte JMN. Hormone-sensitive lipase is localized at synapses and is necessary for normal memory functioning in mice. J Lipid Res. 2022 May;63(5):100195. doi: 10.1016/j.jlr.2022.100195. Epub 2022 Mar 15. PMID: 35300984; PMCID: PMC9026619.

52. Skoug C, Rogova O, Spégel P, Holm C, Duarte JMN. Genetic deletion of hormone-sensitive lipase in mice reduces cerebral blood flow but does not aggravate the impact of diet-induced obesity on memory. J Neurochem. Published online February 5, 2024. doi:10.1111/jnc.16064

53. Slomnicki LP, Pietrzak M, Vashishta A, Jones J, Lynch N, Elliot S, Poulos E, Malicote D, Morris BE, Hallgren J, Hetman M. Requirement of Neuronal Ribosome Synthesis for Growth and Maintenance of the Dendritic Tree. J Biol Chem. 2016 Mar 11;291(11):5721–5739. doi: 10.1074/jbc.M115.682161. Epub 2016 Jan 12. PMID: 26757818; PMCID: PMC4786710.

54. Soares AF, Duarte JMN, Gruetter R. Increased hepatic fatty acid polyunsaturation precedes ectopic lipid deposition in the liver in adaptation to high-fat diets in mice. MAGMA. 2018 Apr;31(2):341–354. doi: 10.1007/s10334-017-0654-8. Epub 2017 Oct 12. PMID: 29027041.

55. Thaler JP, Yi CX, Schur EA, Guyenet SJ, Hwang BH, Dietrich MO, et al. (2012). Obesity is associated with hypothalamic injury in rodents and humans. J. Clin. Invest. 122, 153–162. doi: 10.1172/JCI59660.

56. Tracy, L.M., Bergqvist, F., Ivanova, E.V., Jacobsen, K.T., Iverfeldt, K. (2013). Exposure to the saturated free fatty acid palmitate alters BV-2 microglia inflammatory response. Journal of molecular neuroscience. 51(3):805–12. doi: 10.1007/s12031-013-0068-7.

57. Tsutsumi R, Hori Y, Seki T, Kurauchi Y, Sato M, Oshima M, Hisatsune A, Katsuki H. (2019). Involvement of exosomes in dopaminergic neurodegeneration by microglial activation in midbrain slice cultures. Biochem Biophys Res Commun. 511(2):427–433. doi: 10.1016/j.bbrc.2019.02.076.

58. Ulloth JE, Casiano CA, De Leon M. Palmitic and stearic fatty acids induce caspase-dependent and -independent cell death in nerve growth factor differentiated PC12 cells. J Neurochem. 2003 Feb;84(4):655–68. doi: 10.1046/j.1471-4159.2003.01571.x. PMID: 12562510; PMCID: PMC4157900.

59. Valdearcos, M., Douglass, J.D., Robblee, M.M., Dorfman, M.D., Stifler, D.R., Bennett, M.L., Gerritse, I., Fasnacht, R., Barres, B.A., Thaler, J.P., Koliwad, S.K. (2017). Microglial Inflammatory Signaling Orchestrates the Hypothalamic Immune Response to Dietary Excess and Mediates Obesity Susceptibility. Cell Metab. 26(1):185–197.e3. doi: 10.1016/j.cmet.2017.05.015.

60. Valdearcos, M., Robblee, M.M., Benjamin, D.I., Nomura, D.K., Xu, A.W., Koliwad, S.K. (2014) Microglia dictate the impact of saturated fat consumption on hypothalamic inflammation and neuronal function. Cell Rep. 9, 2124–2138. doi: 10.1016/j.celrep.2014.11.018.

61. Wang, L., Pavlou, S., Du, X., Bhuckory, M., Xu, H., Chen, M. (2019). Glucose transporter 1 critically controls microglial activation through facilitating glycolysis. Mol Neurodegener (2019) 14(1):2. 10.1186/s13024-019-0305-9

62. Wiklander OPB, Brennan MÁ, Lötvall J, Breakefield XO, El Andaloussi S. (2019). Advances in therapeutic applications of extracellular vesicles. Sci Transl Med. 15;11(492):eaav8521. doi: 10.1126/scitranslmed.aav8521.

63. Winston CN, Romero HK, Ellisman M, Nauss S, Julovich DA, Conger T, Hall JR, Campana W, O’Bryant SE, Nievergelt CM, Baker DG, Risbrough VB, Rissman RA. (2019). Assessing Neuronal and Astrocyte Derived Exosomes From Individuals With Mild Traumatic Brain Injury for Markers of Neurodegeneration and Cytotoxic Activity. Front Neurosci. 2;13:1005. doi: 10.3389/fnins.2019.01005.

64. Won, S. J., Yoo, B. H., Kauppinen, T. M., Choi, B. Y., Kim, J. H., Jang, B. G., … Suh, S. W. (2012). Recurrent/moderate hypoglycemia induces hippocampal dendritic injury, microglial activation, and cognitive impairment in diabetic rats. Journal of Neuroinflammation, 9:182. doi: 10.1186/1742-2094-9-182.

65. Yang Y, Boza-Serrano A, Dunning CJR, Clausen BH, Lambertsen KL, Deierborg T. (2018). Inflammation leads to distinct populations of extracellular vesicles from microglia. J Neuroinflammation. 15(1):168. doi: 10.1186/s12974-018-1204-7.

66. Zhang P, Zhou X, He M, Shang Y, Tetlow AL, Godwin AK, Zeng Y. (2019). Ultrasensitive detection of circulating exosomes with a 3D-nanopatterned microfluidic chip. Nat Biomed Eng. 3(6):438–451. doi: 10.1038/s41551-019-0356-9.

